# Genetic perturbation enhances functional heterogeneity in alkaline phosphatase

**DOI:** 10.1101/2021.11.25.470051

**Authors:** Morito Sakuma, Shingo Honda, Hiroshi Ueno, Kentaro Miyazaki, Nobuhiko Tokuriki, Hiroyuki Noji

## Abstract

Enzymes inherently exhibit molecule-to-molecule heterogeneity in catalytic activity or function, which underlies the acquisition of new functions in evolutionary processes. However, correlations between the functional heterogeneity of an enzyme and its multi-functionality or promiscuity remain elusive. In addition, the modulation of functional heterogeneity upon genetic perturbation is currently unexplored. Here, we quantitatively analyzed functional heterogeneity in the wild-type and 69 single-point mutants of *Escherichia coli* alkaline phosphatase (AP) by employing single-molecule assay with a femtoliter reactor array device. Most mutant enzymes exhibited higher functional heterogeneity than the wild-type enzyme, irrespective of catalytic activity. These results indicated that the wild-type AP minimizes functional heterogeneity, and single-point mutations can significantly expand the span of functional heterogeneity in AP. Moreover, we identified a clear correlation between functional heterogeneity and promiscuous activities. These findings suggest that enzymes can acquire greater functional heterogeneity following marginal genetic perturbations that concomitantly promote catalytic promiscuity.

## Introduction

As our understanding of enzymes matured over the years, the once simple and prevailing ―one enzyme, one function‖ paradigm has undergone a change. Advances in technology and statistical methods have revealed that enzymes intrinsically exhibit molecule-to-molecule heterogeneity in catalytic activity or substrate specificity, even in a clonal population^1–3^. Such functional heterogeneity among molecules persists on time scales far beyond thermal fluctuations - from milliseconds to hours in some enzymes^4–18^. Therefore, it can be referred to as long-lived functional substates; for simplicity of discussion in this work, we define ―functional substate‖ as the long-lived and distinct functional states among the enzyme molecules with the same genotype, independent of any genetic mutation.

The functional substate of enzyme molecules is considered key for molecular mechanisms involving enzymatic functions^4–16^ and may be driving the acquisition of new functions during evolution^17–20^. Specifically, functional substates have been proposed to underlie enzyme promiscuity (or multi-functionality): within an ensemble of molecules, certain dominant functional substates can exclusively perform the native function, whereas some minor functional substates catalyze secondary or promiscuous reactions^3, 21–24^. Moreover, it is postulated that the evolutionary optimization of enzyme function can occur by shifting the population balance among functional substates^3, 21–24^. This view hypothesizes that enzymes with well-optimized native activities have become specialists throughout evolution by converging toward a limited number of functional substates. Conversely, it can be expected that random mutations have the potential to disrupt the functional substates optimized for the native activity, thus to enhance promiscuous functions that were previously supported by a low substate population. Hence, manipulating the functional substate of an enzyme could be key to understanding enzyme evolution and harnessing promiscuous activities for enzyme engineering.

The majority of our understanding regarding functional substates is based on structural studies. Such long-lived functional substates are thought to originate from stable conformational substates revealed by structural analyses using X-ray crystallography, nuclear magnetic resonance (NMR), and molecular dynamics (MD) simulations^25, 26^. Previous studies have revealed that function-altering mutations can perturb the protein structure and increase conformational heterogeneity^27, 28^. However, it is unclear whether such conformational heterogeneity results in functional substates. Moreover, the averaging of a population of enzymes in crystallography or NMR would mean subtle conformational changes in enzymes, even in stable and distinct populations of functional substates, barely detectable by the structural analysis and hence remain obscured^29, 30^. Therefore, direct experimental measurement of functional substates that link each enzyme molecule with its functional characteristics is essential to advance our understanding.

The development of single-molecule enzymology techniques using microreactor/microfluidic devices and microscopic measurements has revolutionized the study of enzymatic reactions^10–16, 31–37^. Recent studies, which utilized a femtoliter reactor array device (FRAD) or similar technologies, statistically analyzed the functional heterogeneity in an enzyme population performing thousands of single-molecule reactions conducted in parallel^14–16, 34–37^. The single-molecule assay can thus provide significant information to determine the functional substate, such as a number of activity states or the extent of the functional heterogeneity, *i.e.*, width of the distribution in catalytic activities. The existence of functional substates observed over time scales of tens of minutes or hours was reported for β-galactosidase (GAL), β-glucuronidase (GUS), and alkaline phosphatase (AP)^14–16, 34, 35^. Those enzymes show one or two distinct population in the functional heterogeneity. Interestingly, wild-type GUS exhibits a relatively low heterogeneity (narrow distribution), whereas the mutants of GUS having high catalytic activity toward promiscuous substrates exhibit higher heterogeneity (broader distribution) than the wild-type^34^. However, answers to many of the fundamental questions concerning functional substates remain elusive. Has evolution enriched specific functional substates during the functional optimization of natural enzymes? How easily can mutations modulate functional substates? Are functional substates of a particular native-like substrate associated with promiscuous activity?

In this study, we employed single-molecule technology using the FRAD to measure catalytic activity of the wild-type and 69 mutants of *Escherichia coli* (*E.coli*) AP. We then analyzed their functional substates based on the functional heterogeneity. We found that the wild-type AP (AP-wt) exhibited the lowest functional heterogeneity, and most mutants presented greater functional heterogeneity than the AP-wt. Hence, a single-point mutation in the AP-wt significantly expands its functional substates. We also tested potential correlations involving functional heterogeneity and promiscuity by measuring the catalytic activity of mutants displaying different extent of functional heterogeneity against various promiscuous substrates. Mutants exhibiting high functional heterogeneity also showed high promiscuous activities, which suggests that the expansion of the functional substates of native function could confer catalytic promiscuity. This study provides the first quantitative evidence of the contribution of molecule-to-molecule heterogeneity for enzyme promiscuity and its evolution.

## Results

### Single-molecule assay of *E. coli* AP

We employed *E. coli* AP as a model to characterize heterogeneity in catalytic activities. AP is a homodimeric metalloenzyme encompassing two Zn^2+^ ions and one Mg^2+^ in its active site (**Fig. 1a**). AP was expressed using the PURE system, a reconstituted cell-free protein synthesis system optimized in our previous studies^16, 36^. The catalytic activity of AP expressed in the PURE system can be measured using a FRAD without purification^16^. After AP molecules were expressed, the PURE reaction solution was diluted and mixed with a fluorescein-based fluorogenic phosphate monoesters substrate (fluorescein diphosphate, FDP) (**Fig. 1a**). Subsequently, to measure single enzyme activity, the enzyme assay mixture was applied to a flow channel of the FRAD displaying over 46,000 reactors (**Fig. 1a** and **Supplementary Fig.1**). A fluorinated oil (FC-40) was then injected into the flow channel to displace the enzyme assay mixture and seal the reactors (**Supplementary Fig.1b**), ensuring negligible cross-reactor diffusion levels over the course of the experiment (**Supplementary Fig. 2**). The single enzyme activity was measured by the increase in the fluorescence intensity in the reactors (**Fig. 1b**). The single-molecule assay was performed according to previously optimized reaction conditions using 1 mM FDP, since AP exhibits catalytic activity even after 6 h without any significant decrease in activity^16^.

**Figure 1.**
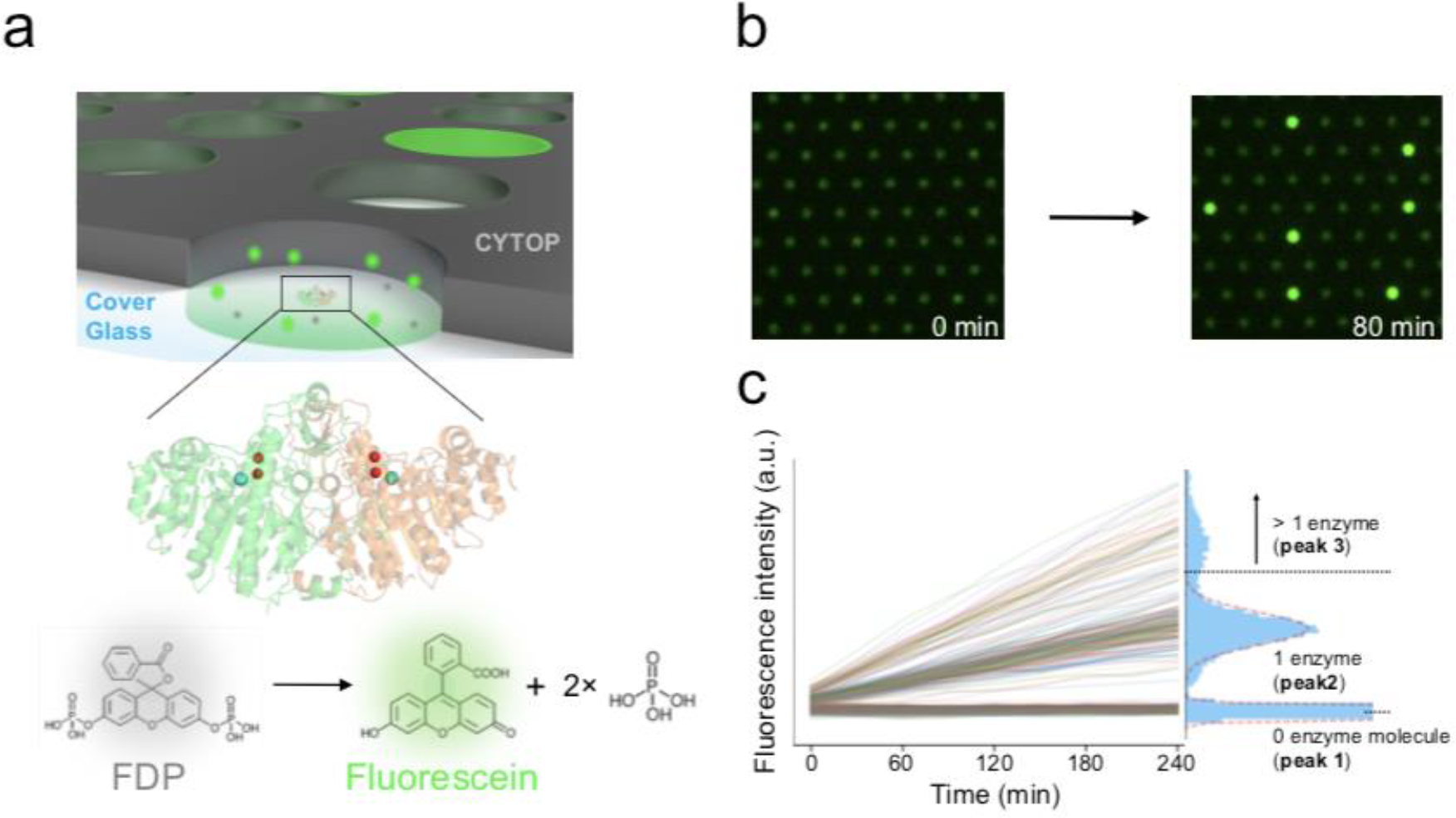
FRAD for measuring single enzyme activity. (**a**) Schematic image of single-molecule assay in the FRAD. In each well or ―reactor‖ patterned on amorphous perfluoro polymer (CYTOP), a single AP molecule hydrolyzes non-fluorescent FDP substrate molecules to release fluorescent product molecules (fluorescein), which accumulates in the reactor. Red and magenta spheres in the crystal structure of dimeric AP-wt (PDB ID: 1ED8) represent Zn^2+^ and Mg^2+^ ions in the active site. (**b**) Snapshots of time-course measurements of catalytic activities observed by microscopy. Reactors containing AP-wt molecules exhibit increase in fluorescence intensity. (**c**) Typical time-course measurement of fluorescence intensity in reactors. The histogram shows the distribution of fluorescence intensity at 240 min. The histogram typically presents three peaks: peak 1 (enzyme-free reactors), peak 2 (single enzyme activity), and peak 3 (containing two or more enzyme molecules in a single reactor). The dotted red lines indicate fitting with Gaussian functions.

### Measurement of heterogeneity in catalytic activities

The catalytic activity and heterogeneity in catalytic activities among AP-wt molecules were evaluated based on the distribution of fluorescence intensity observed in all reactors (**Fig. 1c**). The histograms of fluorescence intensity typically showed two main discrete peaks: a peak of low fluorescence intensity was derived from enzyme-free reactors (peak 1) and another (peak 2) with higher fluorescence intensity represents reactors containing a single enzyme molecule. We occasionally observed an additional minor peak (peak 3) containing two or more enzyme molecules, which exhibited approximately 2-fold higher intensity than peak 2^16, 35^.

The enzyme solution was diluted in order to reduce the population of peak 3 reactors, and if peak 3 remained, it was excluded from the analysis. We measured heterogeneity in catalytic activities by fitting the distribution of peaks 1 and 2 with a Gaussian function to determine the mean intensity (*MI*_1 or 2_) and standard deviation (*SD*_1 or 2_) of the peaks (see **Methods**). Using the *MI* of the two peaks and measurement time (*t*), the mean catalytic activity (*MA*) across all single enzyme molecules is defined as:

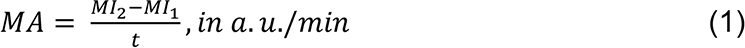

The *MA* of the AP-wt with 1 mM FDP was determined to be 13.2 ± 0.8 (a.u./min) (**Extended Data**), which corresponds to the turnover rate of 19 s^-1^ calculated using the calibration curve (**Supplementary Fig. 4a**), consistent with the turnover rate determined in bulk measurement, 26 s^-1^ (**Supplementary Fig. 3b and 4b**). The concentration of FDP in the single-molecule assay was sufficiently higher than the *K*_m_ obtained in bulk measurements (*K*_m_ = 8 µM) (**Supplementary Fig. 3a and b**). Heterogeneity in catalytic activities was defined as the coefficient of variation (*CV*_i_ = *SD*_i_/*MI*_i_, i = 1 or 2). The *CV* of AP-wt (*CV*_2_: *CV* for peak 2 = 12 ± 3% in nine independent measurements) was approximately two-fold higher than the inherent noise of the system estimated from the *CV* of peak 1 (*CV*_1_ = 6 ± 1%) (see **Methods**). Finally, we calculated the ―functional heterogeneity‖ of AP-wt as the normalized *CV* (*CV*_n_) by compensating for system noise, *CV*_1_, as:

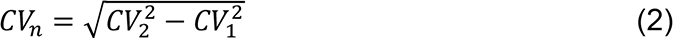

The resulting *CV*_n_ for AP-wt was 10 ± 3%.

### Functional heterogeneity of single-point mutants

To investigate whether the heterogeneity in catalytic activities can be modulated by mutations, we randomly selected 65 single-point mutants of *E. coli* AP from a mutagenized library that we previously generated^36^. In addition, we prepared four mutants (D101S, G118D, D153S, and K328R) that were identified as highly active against FDP (see **Methods**). In total, 69 mutants were examined by single-molecule assay with FRAD. The activity of each mutant was measured in duplicate, and *MA* (a.u./min) and functional heterogeneity (*CV*_n,_ %) were analyzed. Variations in the distribution were observed, as the single enzyme peak (peak 2) could represent a single peak or comprise two neighboring peaks, for example, E66V and D128V (**Supplementary Fig. 5**). A double-Gaussian function could not explicitly fit the neighboring two peaks in some mutants, and thus, all distributions were fitted with a single Gaussian function. The replicates for all mutants exhibited good reproducibility in determining *MA* and *CV*_n_ (**Supplementary Fig. 6** and **Extended Data**).

The four high-activity mutant variants, D101S, G118D, D153S, and K328R — all located around the catalytic reaction center (within 10 Å from the ligand) — exhibited 4 to 13-fold higher activity than AP-wt (**Fig. 2a**). Most randomly chosen mutations only marginally affected *MA*; almost all mutants presented an *MA* within 2- fold of the wild-type (**Fig. 2a**). Furthermore, the 65 randomly selected mutants did not reveal any clear correlation between catalytic activity and the distance of the mutated site from the catalytic reaction center (**Fig. 2a** and **b**).

**Figure 2.**
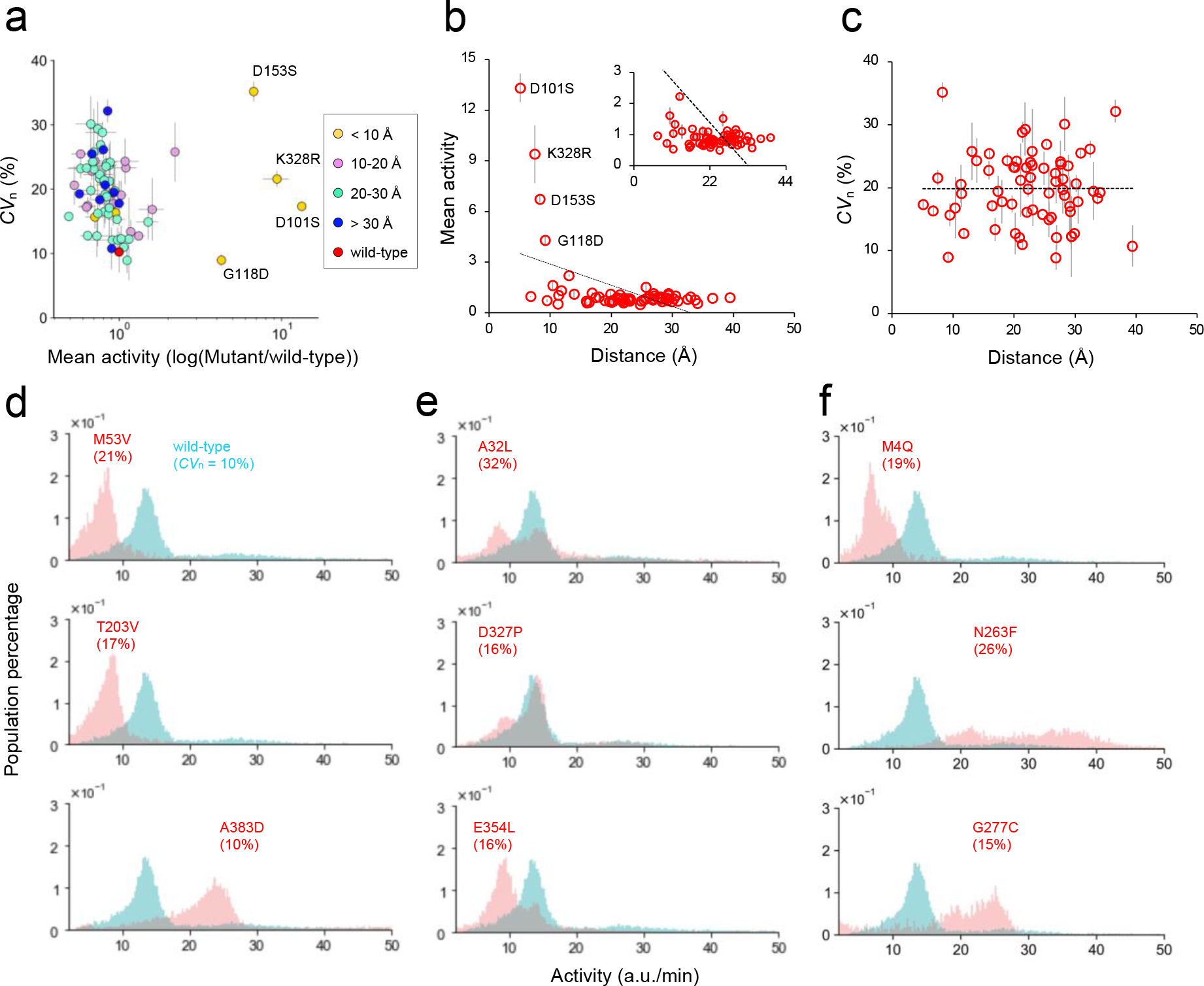
Alteration in catalytic activity and functional heterogeneity by single- point mutations. (**a**) Mean catalytic activity (*MA*) (a.u./min) and functional heterogeneity (*CV*_n_) (%) of AP-wt and mutant APs were plotted (means ± *SD*). The activity of the mutants was normalized to that of the AP-wt, and the x-axis was converted to a logarithmic scale. The data points are colored according to the distance (Å) between a given mutation and the ligand (phosphate) present in the active site to the crystal structure of AP. The red-filled circle represents the AP-wt. (**b**) Dependence of activity on distance from the ligand to mutation sites. The dotted line shows a linear fitting of the plot (slope = -0.13, R^2^ = 0.24). (**c**) Dependence of *CV*_n_ on the distance from ligand to mutation sites. The dotted line shows a linear fitting of the plot (slope = -0.01 × 10^-1^, R^2^ = 0.05 × 10^-4^). (**d**)-(**f**) Representative distributions of mutants that exhibited single peaks shifted from the AP-wt (**d**), additional peaks partially overlapping with the AP-wt (**e**), and shifted additional peaks (**f**)

We found that the AP-wt exhibited a single narrow distribution peak (**Supplementary Fig. 5**); its *CV*_n_ being the lowest among all mutants (*CV*_n_ = 10%). Most mutants showed significantly higher heterogeneity than the AP-wt. On average, the *CV*_n_ of mutants was 2-fold higher (20%), and some mutants showed up to 3-fold higher *CV*_n_ than the AP-wt. Intriguingly, there was no apparent correlation between the *CV*_n_ and *MA* (**Fig. 2a**). This observation suggests that functional heterogeneity is principally independent of the level of catalytic activity and that the molecular mechanism for gaining functional heterogeneity differs from that for catalysis enhancement.

Across all mutants, we observed a broad range of combinations between functional heterogeneity and catalytic activity (**Fig. 2d-f and Supplementary Fig. 7**). Indeed, some mutants exhibited a single peak, yet displayed catalytic activities different from the wild-type (**Fig. 2d**). We hypothesized that, in these specific cases, the mutation likely modulates the transition states of the kinetic bottleneck steps without disturbing the populations of functional substates. By contrast, in other mutants, one population exhibited catalytic activity comparable to that of the AP-wt (**Fig. 2e**), and an additional population was also found, resulting in a bimodal distribution. Mutations that lead to shifts between sub-populations, visible through the activity change in the ensemble average of enzymes, can underlie the emergence of a new functional substate within a clonal population. Such mutations may act to stabilize functional substates, which were minor in the AP-wt. Finally, another set of mutations resulted in two peaks, although both peaks were shifted from that of the AP-wt population (**Fig. 2f**), suggesting that these mutations can simultaneously modulate enzyme kinetics and the population balance between various functional substates. Taken together, these observations suggest that a mutation can easily alter the balance of AP functional substates, and the effect on the population balance differs widely, depending on the specific mutation.

The four highly active AP mutants (D101S, G118D, D153S, and K328R) also exhibited distinct differences in terms of heterogeneity in catalytic activities (**Fig. 3**). D101S, D153S, and K328R exhibited shifted and broader distributions than the AP- wt. Especially, D153S showed the highest *CV*_n_ among the mutants. However, G118D did not enhance heterogeneity despite having four-fold higher catalytic activity than the AP-wt, *i.e.*, the *CV*_n_ of the peak remained identical to the AP-wt but its *MA* was different. Thus, similar to the single-point mutants, wide variation in the heterogeneity in catalytic activities was observed. This high heterogeneity in the highly active AP mutants might be due to structural heterogeneity involving the active site, such as the flipping of catalytically important residues or a large change in phosphate coordination as demonstrated by previous studies^38–41^. In the future, further elucidation of the direct correlation between structural changes in the active site and the alteration of functional substates is pivotal to our understanding of enzyme mechanism and evolution.

**Figure 3.**
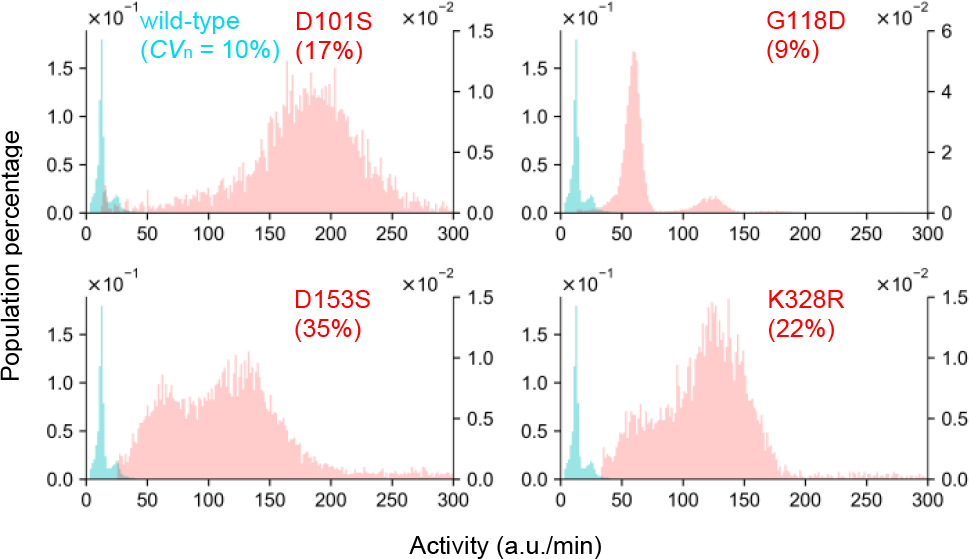
Alteration of catalytic activity and functional heterogeneity in highly active mutants. The distribution of catalytic activity of AP-wt and highly active AP mutants (D101S, G118D, D153S, and K328R). Blue and red bins show the distribution of the AP-wt (first y-axis) and mutants (second y-axis), respectively.

### Functional heterogeneity and structural position of mutations

In order to investigate whether AP possesses mutational hot spots that may be responsible for enhanced activity and/or high functional heterogeneity, we plotted the *MA* and *CV*_n_ of each single mutation against the distance of Cα of mutation sites from phosphate in active site **(Fig. 2b and c**). We also presented a heat map of the crystal structure of AP-wt in order to map mutation sites with higher *MA* and *CV*_n_ on the 3D structure (**Supplementary Fig. 8**). Interestingly, the mutations giving rise to highly active mutants were located in the vicinity of the active site; however, there was no significant correlation between the *CV*_n_ and the distance from the active site. We also analyzed mutations based on their proximity to the two disulfide bonds (C168-C178 and C286-C336) or the interface between the monomers (**Supplementary Fig. 9**)^42, 43^, but there were no clear correlations. Thus, these findings suggest that functional heterogeneity can potentially be modulated globally across the whole enzyme structure.

### Functional heterogeneity and promiscuity

It has been suggested that the expansion of functional substates may be related to the enhancement of reactivity to promiscuous substrates^3, 17–20, 22, 24, 34^. To test this, we measured the activity of AP-wt and four highly active mutants (D101S, G118D, D153S, and K328R) against promiscuous substrates: phosphate diesters (bis-*p*-nitrophenyl phosphate, *bp*NPP) and phosphonate monoesters (*p*-nitrophenyl phenyl phosphonate, *p*NPPP), in addition to another phosphate monoesters substrate, *p*-nitrophenyl phosphate (*p*NPP) (**Fig. 4a**). Then, we analyzed the correlation between *CV*_n_ measured using FDP and the promiscuous activities. *bp*NPP and *p*NPPP can be employed to evaluate the promiscuous activity of AP-wt^44, 45^. Since single-point mutations involving the active site can significantly alter promiscuous activity, including *E. coli* AP^46–48^, those highly active mutants could be used to assess the effect of *CV*_n_ levels (9-35%) on promiscuity. The activities against *bp*NPP and *p*NPPP were too low to be measured at the single-molecule level; hence, we measured their promiscuous activities in a bulk solution with purified enzymes (**Supplementary Fig. 3c**). Although the four AP mutants exhibited higher activity against *p*NPP than the AP-wt (**Fig. 4b**), no correlation was observed between catalytic activity and *CV*_n_ (**Fig. 4c**, *R*^2^ = 0.28), as previously observed with FDP.

**Figure 4.**
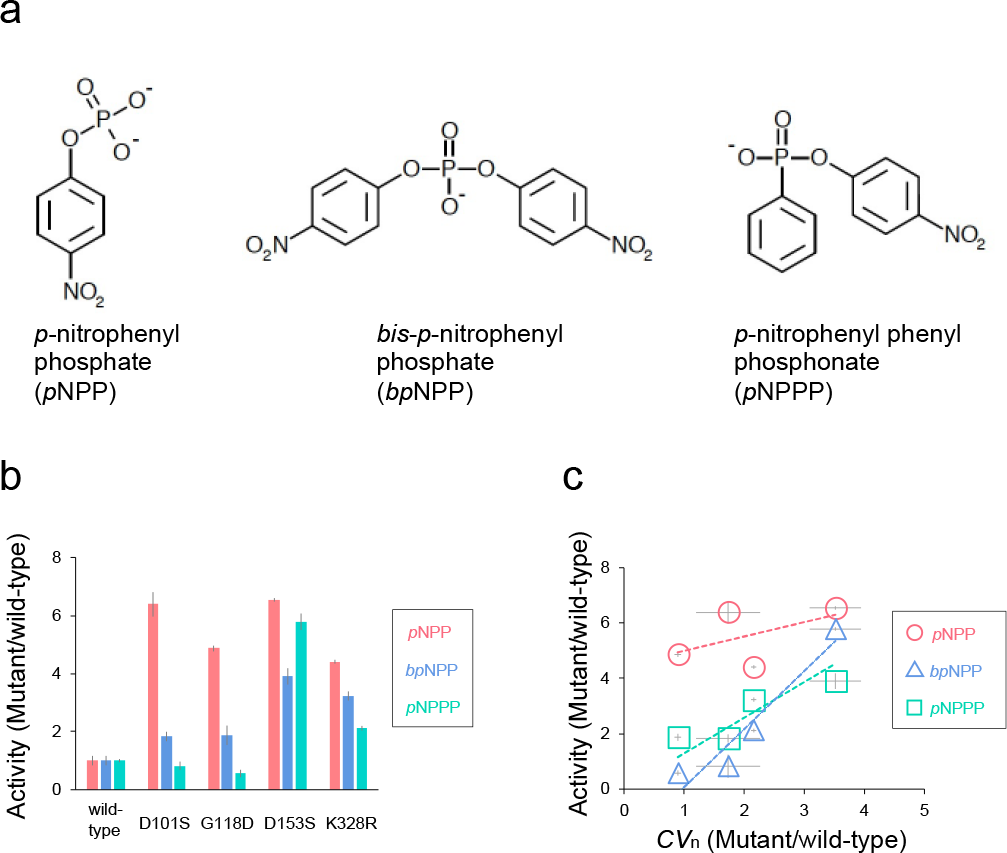
Correlation between functional heterogeneity and catalytic activity against promiscuous substrates. (**a**) Chemical structures of phosphate monoesters (*p*-nitrophenyl phosphate, *p*NPP), phosphate diesters (bis-*p*-nitrophenyl phosphate, *bp*NPP), and phosphonate monoesters (*p*-nitrophenyl phenyl phosphonate, *p*NPPP) substrates. (**b**) Comparative activity of the AP-wt and highly active mutant APs (D101S, G118D, D153S, and K328R) against the substrates in three independent measurements (means ± *SD*) in the bulk solution. The activity of mutants was normalized to that of the AP-wt. (**c**) Correlation between activities against those substrates and *CV*_n_ of the highly active mutants. Activity and *CV*_n_ of mutant AP were comparable to those of AP-wt. The *CV*_n_ of the mutants was measured using the FDP shown in Fig. 3. The slopes of *p*NPP, *bp*NPP, and *p*NPPP were 0.51 (R^2^ = 0.28), 2.10 (R^2^ = 0.91), and 0.85 (R^2^ =0.82), respectively.

Interestingly, however, D153S and K328R, which have higher *CV*_n_, exhibited greater activity than the AP-wt against promiscuous substrates (**Fig. 4b)**. In particular, D153S, which displayed the highest *CV*_n_ among the mutants, showed 4- and 6-fold higher activity against *bp*NPP and *p*NPPP than the AP-wt. On the other hand, D101S and G118D, whose *CV*_n_s were close to AP-wt, showed only modest activity enhancement. To investigate a possible correlation between functional heterogeneity and promiscuity, the activities of the mutants toward promiscuous substrates were plotted against the *CV*_n_ of the mutants (**Fig. 4c**). Clear correlations were found for activity against *bp*NPP (*R*^2^ = 0.91) and *p*NPPP (*R*^2^ =0.82) (**Fig. 4c**). These observations suggested that functional heterogeneity is associated with AP enzyme promiscuity.

## Discussion

In the present study, we assessed the functional heterogeneity of a model enzyme using AP-wt and 69 mutants by performing single-molecule assays with the FRAD. AP-wt showed the lowest level of functional heterogeneity, and most of the mutants showed more significant heterogeneity than the AP-wt. These observations showed that even a single-point mutation in AP-wt is prone to enhancing functional heterogeneity, suggesting that AP-wt is perhaps optimized to reduce functional heterogeneity. A possible scenario is that AP has evolved as a specialist enzyme, minimizing the functional substate toward its native function of phosphate monoesterase. Native enzymes have undergone a long evolutionary process under various selection pressures from diverse environmental conditions; thus, these natural enzymes may have undergone optimization in terms of their conformational and functional substates^21–24^. If the functional substates of AP-wt for its native function have been evolutionary optimized over billions of years, mutations may readily disrupt the fine-tune balance of the substates. As a results, the expansion of the functional substates may lead to high functional heterogeneity in most mutants independent of catalytic activity toward FDP or *p*NPP (**Fig. 3** and **4c**).

Another significant finding in the present study is the clear correlation between functional heterogeneity and promiscuity (**Fig. 4b and c**). AP-wt is highly optimized for the hydrolysis of phosphate monoesters so that AP-wt has low promiscuous activity, >10^9^ lower than the native AP activity^44, 45^, against non-native substrates with phosphate diesters or phosphonate monoesters. Interestingly, an active mutant, D153S, exhibited higher functional heterogeneity and remarkable enhancements of catalytic activity against non-native substrates with *bp*NPP and *p*NPPP than the AP- wt. Similarly, evolved GUS with high promiscuous activity against several glycosidic substrates exhibited more extensive heterogeneity in catalytic activity than the wild-type enzyme^34^. Moreover, wild-type GAL showed higher heterogeneity in catalytic activity than wild-type GUS because it retains three catalytic functions^34, 49^. These findings and our results reinforce the long-discussed concept of enzyme evolution; enzymes can enhance their promiscuity or multi-functionality by gaining functional heterogeneity with multiple functional substates. However, only a handful of enzymes and their variants have been measured at the level of single enzyme kinetics to date, and it is difficult to generalize these observations to other systems. For further generalization of this scenario, more comprehensive studies involving the various types of enzymes are required.

## Materials and Methods

### Creation and cloning of AP for single-molecule assay

The 65 single-point mutants of *E. coli* APs were chosen from a site-directed saturation library mutagenized at single sites (residues 1-449) used in our previous study^36^. Each site was mutagenized using NNK degenerate codons giving rise to 450 individual libraries (one for each position), then subcloned into a T7 expression vector with an ampicillin selective marker (pRSET-B) (Invitrogen^TM^, Thermo Fisher Scientific, Waltham, MA, USA) by Gibson assembly. The mutants used in the single- molecule assays were randomly selected from the library. The plasmid library for each selected position was transformed into HIT-JM109 competent cells (RBC Bioscience, Birmingham, U.K.), and the cells were plated on LB agar plates containing 100 µg/mL ampicillin. After culturing the cells overnight, three to ten colonies were picked for each selected position and cultured separately in the LB media overnight. The plasmids were then purified from the cultured cells and mutation sites were confirmed by DNA sequencing. For each mutation position, one to eight plasmids were obtained using this method, and one plasmid was randomly selected from those plasmids and used for expression with a PURE system (PURExpress In Vitro Protein Synthesis Kit, New England BioLabs, Ipswich, MA, USA).

Of the highly active mutant APs, D101S was kindly gifted by Abbott Laboratories^38^. The three remaining highly active mutants (G118D, D153S, and K328R) were obtained by performing a directed evolution starting from the library described above and a random mutagenesis library constructed by error-prone PCR (Agilent, Santa Clara, CA, USA). We used a FRAD to perform directed evolution under identical conditions to our previous study^36^. As the previous 65 variants, these four mutants were also subcloned into the pRSET-B vector for expression.

### Expression of AP for the single-molecule assay

For single-molecule assays, wild-type and mutant APs were expressed using PURE system. The PURE system includes all proteins required for transcription and translation, and proteins can be expressed by mixing the PURE system and DNA that encodes the sequence of proteins downstream of a T7 promoter. To express the APs, PURE components A and B, disulfide bonding enhancer 1 and 2 (New England BioLabs), RNase inhibitor (Takara Bio Inc., Shiga, Japan), 100 µM ZnCl_2_, and ∼50 ng of the pRSET-B expression vector were mixed by vortexing and incubated at 27°C for 4 h in a thermocycler. Supplementation of disulfide bond enhancers and ZnCl_2_ promotes dimer formation of fully activated APs without loss of their bonds and zinc ions^16^. Note that the levels of endogenous AP in the PURE system was negligible, and thus, the expressed AP could be directly used for measurements without purification after dilution to an optimum concentration^16^.

### Expression and purification of AP for bulk reaction

For bulk reactions, a vector (pET-21(+)) for large-scale expression and purification in *E. coli* was custom-synthesized (Twist Bioscience, South San Francisco, CA, USA). The pET-21(+) vector was designed to have a signal peptide^50^ at the N-terminus and a StrepII-tag on the C-terminus. The sequences of AP-wt, D101S, G118D, D153S, and K328R were inserted between the signal peptide and the StrepII-tag.

AP-wt and highly active mutants were expressed using OverExpress C43(DE3) cells. The pET-21(+) expression vector bearing the AP sequence was transformed into cells by electroporation. The transformed cells were pre-cultured overnight at 37°C in 8 mL of the LB media containing 100 µg/mL ampicillin. Then, 2 mL of the culture solution was transferred to 200 mL of Overnight Express Instant TB medium (MilliporeSigma, Burlington, MA, USA) in a 2 L flask and cultured at 37°C for 8 h, and then the AP protein was expressed at 30°C for 16 h. The cell lysis steps followed a previously established osmotic shock method^41^. The cell lysate passed over a Strep-Tactin column (IBA Lifesciences, Goettingen, Germany) pre-equilibrated with a binding buffer (10 mM Tris-HCl (pH 7.4), 200 mM NaCl, and 10 µM ZnCl_2_) and washed with ten column volumes of the clean binding buffer. Then, proteins were eluted using the elution buffer (20 mM biotin, 10 mM Tris-HCl (pH 7.4), 200 mM NaCl, and 10 µM ZnCl_2_), which was subsequently exchanged to a storage buffer (10 mM sodium MOPS, pH 7.0, 50 mM NaCl, 200 µM ZnCl_2_, and 1 mM MgCl_2_) using a desalting column (Econo-Pac 10DG, BioRad, Hercules, CA, USA). Finally, the protein solutions were concentrated by centrifugation (Spin filters 10k, Pall Laboratory, Port Washington, NY, USA), and their purities were analyzed by SDS-PAGE (MilliporeSigma).

### Fabrication of FRAD

The FRAD was fabricated on a cover glass via photolithography and etching^16, 35–37^. The cover glass (24 × 32 mm, thickness 0.13-0.17 mm, Matsunami Glass Ind., Osaka, Japan) was sonicated in 10 M KOH solution (FUJIFILM Wako Pure Chemical Industries, Osaka, Japan) for 10 min. After washing with deionized water and drying under a nitrogen stream, an amorphous fluorocarbon polymer (CYTOP, Asahi Glass, Tokyo, Japan) was coated onto the glass at 4,000 rpm for 30 s using a spin coater. The glass was then baked at 180°C for 30 min. A positive photoresist (AZ-4903, Merck KGaA, Darmstadt, Germany) was coated onto the CYTOP-coated glass at 7,500 rpm for 60 s. The glass was baked at 55°C for 3 min and at 110°C for 10 min. Photolithography was then performed using a mask aligner (BA100it, Nanometric Technology, Tokyo, Japan) with a chrome photomask with 3 µm holes. The photoresist on the glass was then developed using a photoresist developer (AZ300 MIF, Merck KGaA) for 5 min. The patterned glass coverslip was washed with deionized water. After drying on a hot plate, the glass was etched by O_2_ plasma using a reactive ion etching machine (RIE-10NR, Samco, Santa Clara, CA, USA). Then, the remaining photoresist on the glass was entirely removed using acetone, and the glass was rinsed with 2-propanol and deionized water. The diameter and thickness of the reactor were measured using a 3D laser scanning microscope (VK-X200, KEYENCE, Osaka, Japan). The diameter and thickness of the reactor array used in these experiments were 2.5 - 3.0 µm and 0.45-0.50 µm, respectively.

### Reagents and solutions for the measurement of catalytic activity

To measure the activity of AP in the bulk solution or FRAD, 1 M diethanolamine (DEA) buffer (pH 9.2) (FUJIFILM Wako Pure Chemical Industry) was used as a reaction buffer^16, 35–37^. To maximize enzyme activity, ZnCl_2_ and MgCl_2_ (FUJIFILM Wako Pure Chemical Industry) were added to the reaction buffer (final concentrations of ZnCl_2_ and MgCl_2_ were 100 µM)^35^. A stock solution of FDP (AAT Bioquest, Sunnyvale, CA, USA) in double-distilled water (ddH_2_O), *p*NPP (MilliporeSigma) in ddH_2_O, *bp*NPP (MilliporeSigma) in ddH_2_O, *p*NPPP (MilliporeSigma) in dimethyl sulfoxide (DMSO), and fluorescein dye (FUJIFILM Wako Pure Chemical Industry) in DMSO were stocked at -20°C and only diluted to 1 mM prior to the measurement.

### Single-molecule assays with FRAD

The FRAD was immersed in 2-propanol and sonicated in a water bath for 10 min. After sonication, the reactor array was thoroughly washed with excess ddH_2_O and dried on a hot plate. The flow channel was assembled on the surface of the FRAD with double-sided adhesive tape (TERAOKA SEISAKUSHO, Tokyo, Japan) and a custom-made upper glass, which has inlet and outlet holes for solution and oil flow to the FRAD (**Supplementary Fig. 1a**)^16, 35–37^. The upper glass was coated with CYTOP to reduce the affinity of the surface for the water solution. The double-sided tape was cut out in the shape of the flow path using a cutting machine (STIKA, Roland DG, Shizuoka, Tokyo), and the tape was applied to the surface of the FRAD.

Then, the upper glass was applied to the tape, and the FRAD was degassed for tight binding. The reaction solution (10 µL) was prepared by mixing AP and a fluorogenic substrate (FDP) in the DEA buffer on ice. Then, the reaction solution was added to the flow channel by pipetting (**Supplementary Fig. 1b**), and the air in the reactor was completely degassed by sonication for 1 s in a water bath. Fluorinated oil (FC- 40, Merck, Kenilworth, NJ, USA) was used to remove the excess amounts of the reaction solution and to seal the reactor (**Supplementary Fig. 1b**). Measurements commenced 5 min after the reactor was sealed. New flow channel of FRAD was used for each assay.

### Fluorescence microscopy for single-molecule assay

All images of single-molecule assays were acquired using an inverted fluorescence microscope (Eclipse Ti2, Nikon, Tokyo, Japan) equipped with a 40 × objective lens (Nikon, Plan Apo, NA = 0.9), multi-wavelength LED illumination (X-cite Turbo, Excelitas, Waltham, MA, USA), sCMOS camera (Zyla, Andor Technology, Belfast, UK), an incubator for temperature control (TOKAI HIT Co., Ltd., Shizuoka, Japan), and a Perfect Focus System (Nikon). The fluorescence of the fluorescein dye was measured using filter sets (excitation 480/15 nm, dichroic 505 nm, emission 535/20 nm, Nikon). All measurements were performed using the NIS-Element software (Nikon). In end-point measurements, 90 different fields of view were taken with 100 ms exposure time. The temperature of the incubator was set to 28°C for all measurements.

The leakage of the fluorophore (*i.e.*, the reaction product) from the reactor was evaluated by the photobleaching of a single reactor (**Supplementary Fig. 2**). Photobleaching was performed using a confocal microscope (Eclipse Ti-E microscope, Nikon) equipped with a 60 × objective lens (Plan Apo, NA = 1.40, Nikon) and a blue laser (488 nm, Coherent). Photobleaching and time-course measurements were performed using NIS-Element software.

### Analysis of single-molecule assay data

Fluorescent images of FRAD were analyzed using a custom Python code. Fluorescence intensity histograms were plotted after analyzing the fluorescence intensity in the FRAD (**Fig. 1c**). Enzyme activity was determined as follows: the first peak was fitted by a single Gaussian distribution, and the mean intensity (*MI_1_*) and standard deviation (*SD_1_*) of peak 1 were analyzed. The single enzyme activity was observed at a threshold where the fluorescence intensity was > *MI_1_* + 5 × *SD_1_*. Then, the distribution of single enzyme activity (peak 2) was fitted by a single Gaussian distribution in order to extract the mean intensity (*MI_2_*) and *SD* (*SD_2_*). For all mutants, distinct peak 2 separated from peak 1 was observed after more than 4 h of measurement. The turnover number (*k_cat_* (s^-1^)) was calculated from *MA* by using a calibration curve providing the concentration of the fluorescein as a function of fluorescence intensity in the FRAD (**Supplementary Fig. 4a**).

*CV*_1_ is the lower detection limit of enzyme activity heterogeneity and represents systematic error of variations in the volume of each reactor and optical noise created by the detection system of the microscope. In this experiment, *CV*_2_, the distribution of catalytic activities of all APs, was always > *CV*_1_ (the mean value of *CV*_1_ in all measurements was 7 ± 2%, mean ± *SD*).

In our FRAD, the number of enzyme molecules in a single reactor follows a Poisson distribution^16, 35–37^. Therefore, the probability and expected average number of enzyme molecules per reactor ( ) can be calculated using the Poisson’s equation:

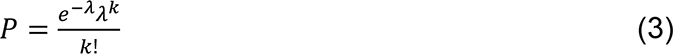

where is the physical number of enzyme molecules in a reactor. can be calculated from the number of positive reactors, including enzyme molecules, and the total number of reactors^16, 35–37^. The mean in all measurements was 0.17 ± 0.13 (mean ± *SD*). With this dilution, we expected that approximately 14% (∼6,400) of the reactors contained the desired single enzyme molecule and ∼2% (∼900) contained two enzyme molecules, and the remaining ∼84% (∼38,600) reactors were empty, that is, without any enzymes.

### Measurement of catalytic activity in bulk solution

AP activity in the bulk solution was measured in a 384-well plate by fluorometry using a plate reader (Synergy H1, BioTek, Winooski, VT, USA). For measurements with FDP, the excitation and emission wavelengths were set to 488 nm and 532 nm, respectively. For the absorbance measurements with *p*NPP, *bp*NPP, and *p*NPPP, the wavelength was set to 405 nm. Changes in fluorescence intensity and absorbance were acquired at two-minute intervals at 28°C. A reaction solution (50 μL) was prepared by mixing AP and substrate in DEA buffer on ice, and then the mixture was applied to the plate wells. The activity was measured after 5 min to ensure temperature equilibration. Three independent measurements were taken to analyze the mean and *SD* of enzyme activity. *p*NPP, *bp*NPP, and *p*NPPP (1 mM) were used to measure the promiscuous activity of APs. The activities of *p*NPP, *bp*NPP, and *p*NPPP were measured for 20 min or 6 h. Enzyme activity was determined from a linear fit of the increase in fluorescence intensity or absorbance over time. The initial rate for FDP was measured using seven different concentrations and fitted by the Michaelis-Menten equation using KaleidaGraph (Synergy Software, Reading, PA, USA).

## Acknowledgements

The authors are grateful to Kenji Akama and Naoki Soga for their technical assistance and to Toshiharu Suzuki and members of the Tokuriki lab for helpful discussions and revision of the manuscript. This work was supported by the ImPACT Program of the Council for Science, Technology, and Innovation, Japan Science and Technology Agency (JST) (to H.N.), Grant-in-Aid for Scientific Research on Innovation Areas from the Japan Society for the Promotion of Science (JSPS) (JP19H05380 to H.U.), Grant-in-Aid for Scientific Research (S) from JSPS (JP19H05624 to H.N.), CREST program from JST, (JPMJCR19S4 to H.N.), and Human Frontier Science Program (HFSP) Program Grant (RGP0054/2020) (to H.N and N.T.).

## Author Contributions

M.S ., H.U., N.T., and H.N. designed the experiments; M.S., S.H., and K.M. performed experiments and analyzed the data; M.S., N.T., and H.N. wrote the paper.

## Competing interests

The authors declare no competing interests.

---

**Supplementary Figure 1.**
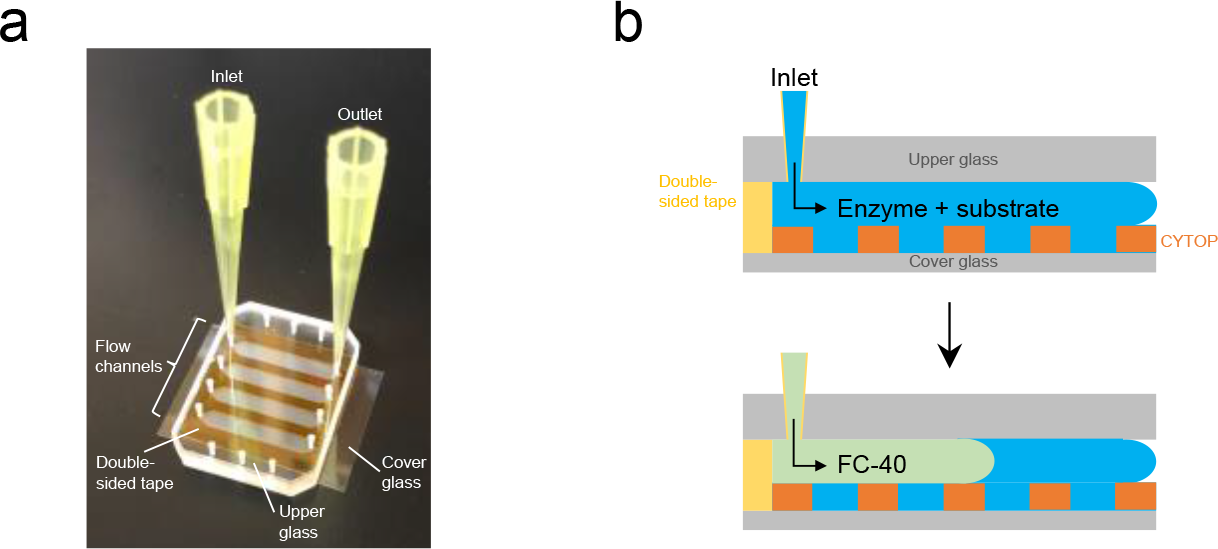
Set up of FRAD for single-molecule assay. (a) The flow channel was assembled on the FRAD. (b) Enzyme assay mixture was injected into the flow channel, and then, FC-40 was applied to displace the assay mixture and seal the reactors.

**Supplementary Figure 2.**
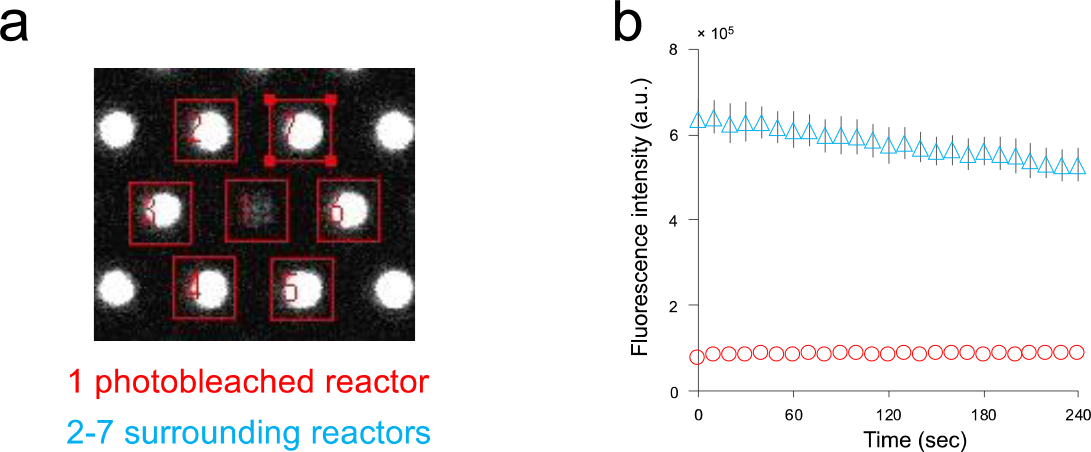
Measuring leakage of fluorescein dye from reactors. (a) One reactor (rectangle 1) was photobleached using a confocal microscope. Changes in fluorescence intensity in photobleached and surrounding reactors (rectangle 2-7) were simultaneously measured. (**b**) Time-course measurement of fluorescence intensity in photobleached and surrounding reactors for 240 min. Red circles indicate fluorescence intensity change of the photobleached reactor; blue triangles represent mean fluorescence intensity and *SD* of the surrounding reactors. The reactors did not show significant fluorescence changes, indicating almost no solution leakage or interchange among the reactors.

**Supplementary Figure 3.**
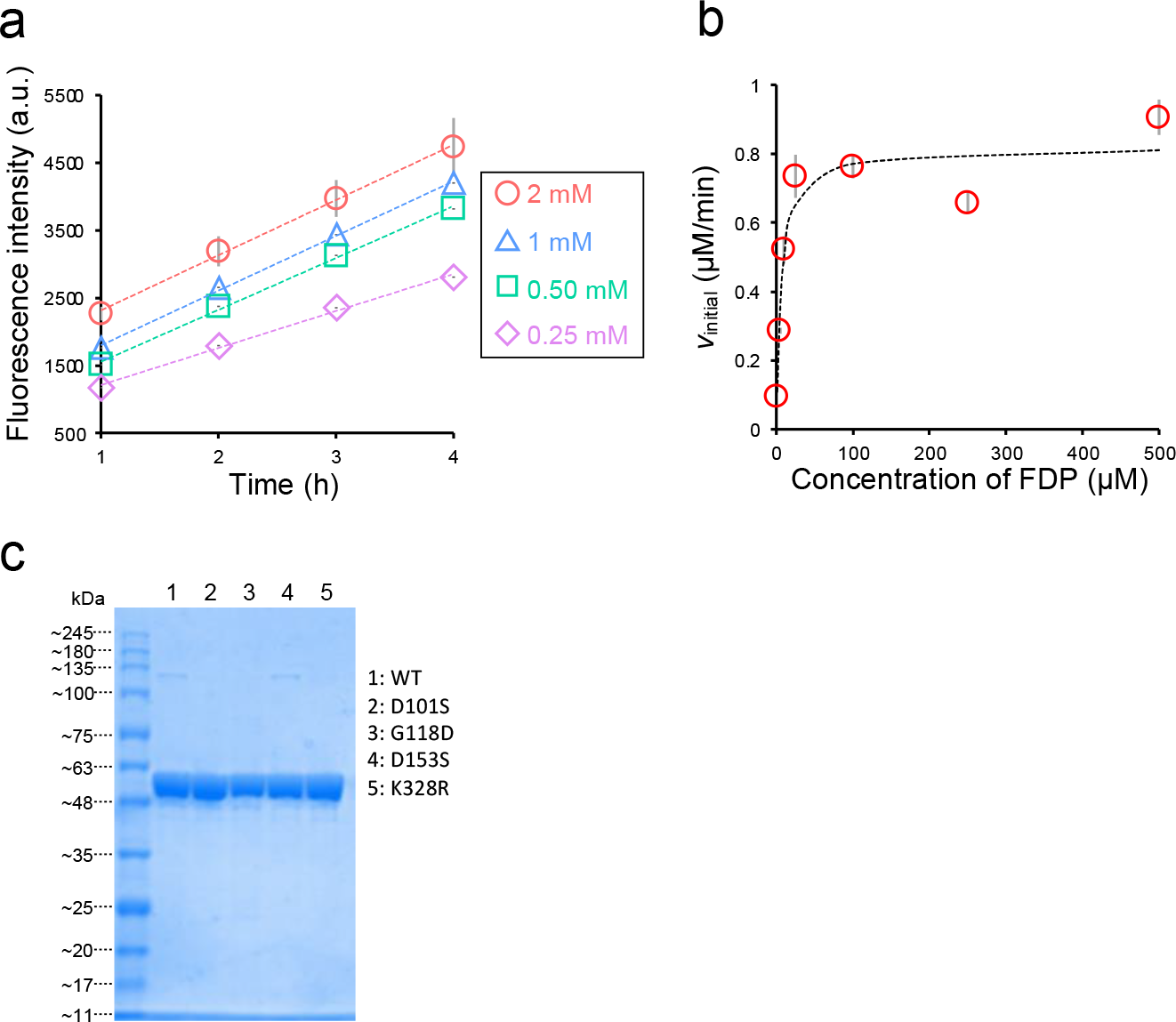
Characterization of catalytic activity of AP-wt in FRAD and bulk solution. (**a**) Time-course measurements of AP-wt in FRAD. The concentrations of FDP were 2 mM (red circles), 1 mM (blue triangles), 0.5 mM (green rectangles), and 0.25 mM (purple diamonds). The plot illustrates changes in mean intensity (*MI_2_*) across three independent measurements (mean ± *SD*). The dotted lines in each plot show a linear fitting of the curves (slopes were 817, 810, 768, and 547 (a.u./h) in 2.0, 1.0, 0.5, and 0.25 mM FDP, respectively, with R^2^ ≧ 0.99). (**b**) Catalytic activity of purified AP-wt was measured in bulk solution with FDP. Red circles show mean initial velocity (*V*_init_, µM/min) and *SD* in three wells. The dotted line shows a fitting with the Michaelis-Menten equation. Analyzed kinetic parameters, *k*_cat_ and *K*_M_, were 27 (s^-1^) and 8 (µM), respectively. (**c**) SDS-PAGE analysis of purified AP-wt and highly active AP mutants. 20 ng of the enzyme was loaded onto the gel. Intense bands were located at the molecular weight of monomeric AP between 48 and 63 kDa corresponding to the monomeric form of AP.

**Supplementary Figure 4.**
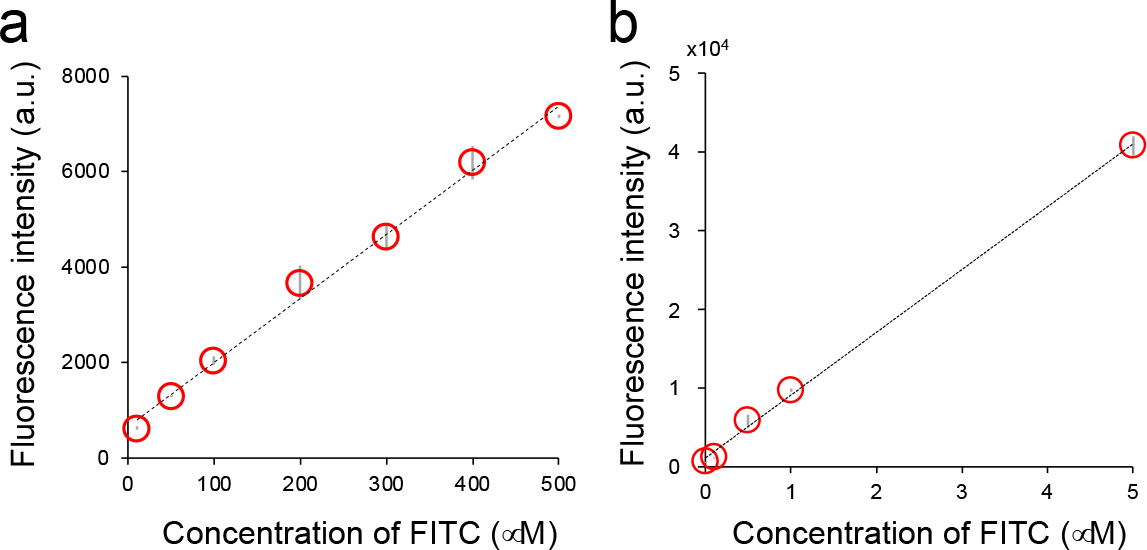
Linear correlation between fluorescein dye concentration and fluorescence intensity in FRAD. (a) and 384-well plates (b). (a) Red circles represent mean intensity and *SD* measured in two different FRADs. Dotted line represents linear fitting of the curves (slope = 13 (a.u./µM), R^2^ = 0.99). (b) Red circles indicate mean intensity and *SD* measured in three different wells. The dotted line represents a linear fitting of the curves (slope = 8,990 (a.u./µM), R^2^ = 0.99).

**Supplementary Figure 5.**
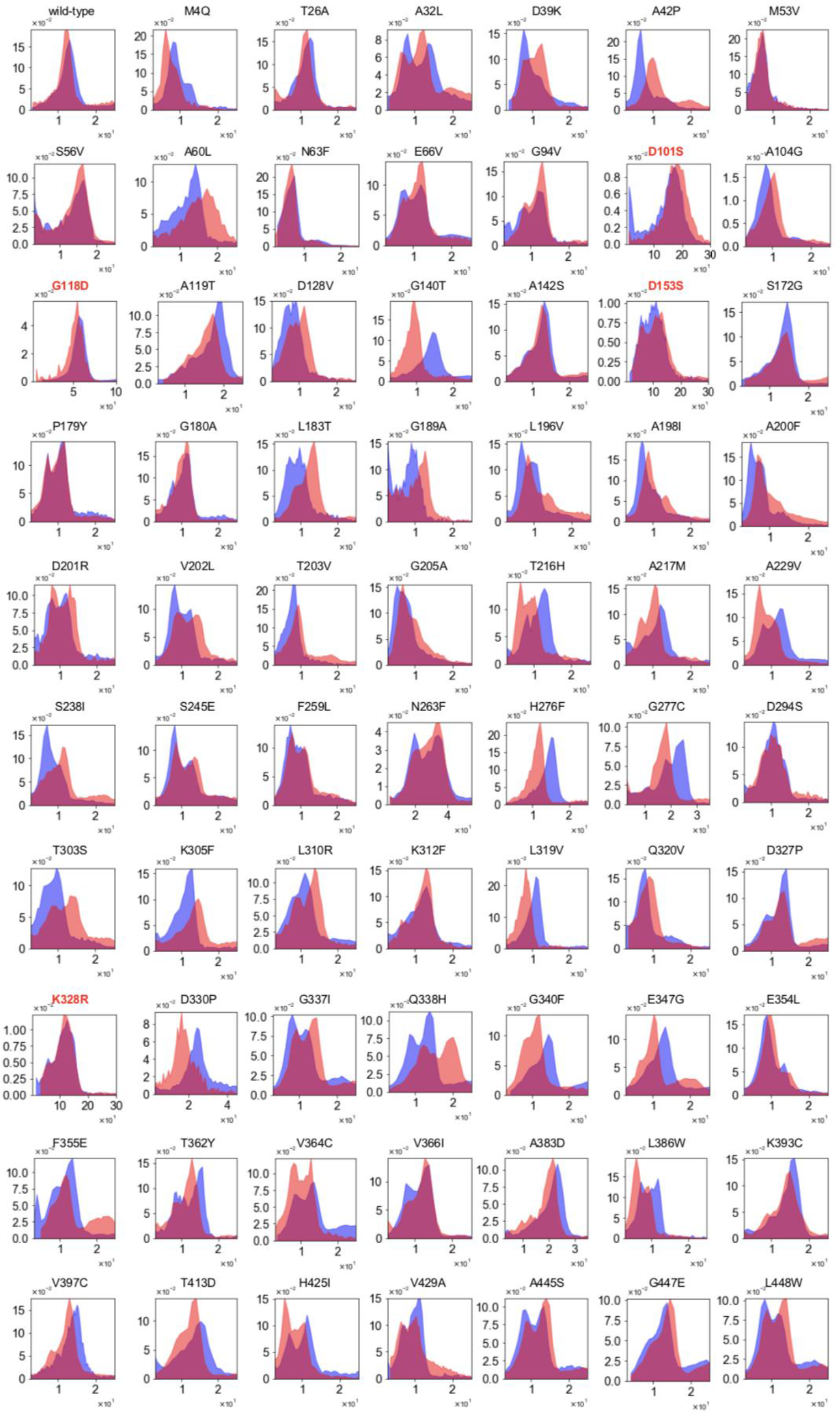
Distribution of catalytic activities measured with FRAD. X- and y-axes represent catalytic activities (a.u./min) and population percentage of enzyme molecules (peaks 2 and 3). Red and blue bins show the distribution of two independent measurements. The identifiers of highly active mutants are shown in red.

**Supplementary Figure 6.**
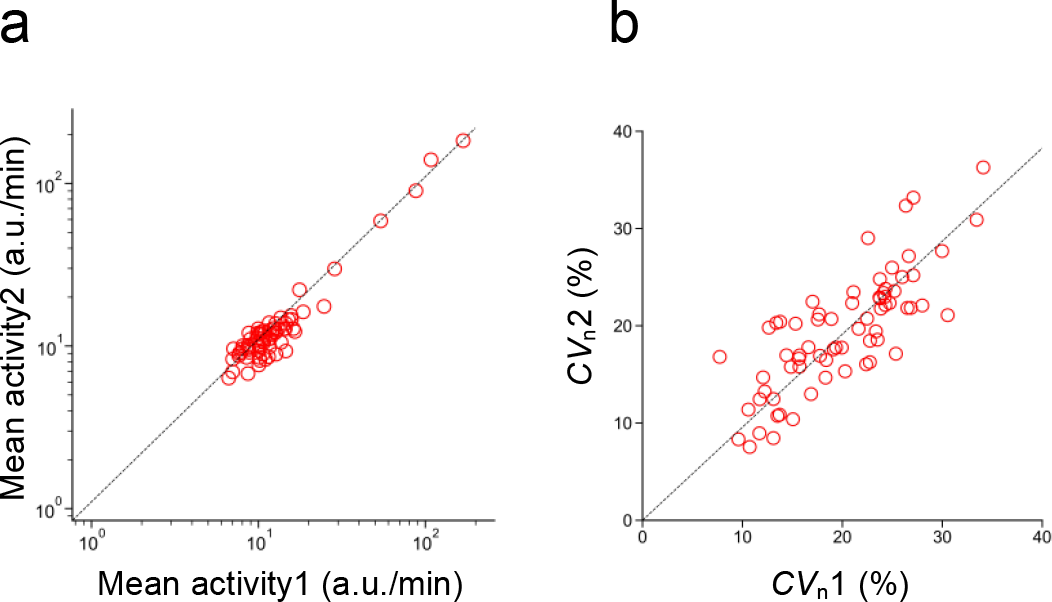
Quality of single-molecule assays with FRAD. (a**) and** (**b**) represent the correlation between two independent measurements of mean catalytic activity (*MA*) and functional heterogeneity (*CV*_n_) of each of the mutants. The dotted lines show linear fittings of the plots. The slopes of (**a**) and (**b**) were 1.1 (R^2^ = 0.99) and 1.0 (R^2^ = 0.97), respectively.

**Supplementary Figure 7.**
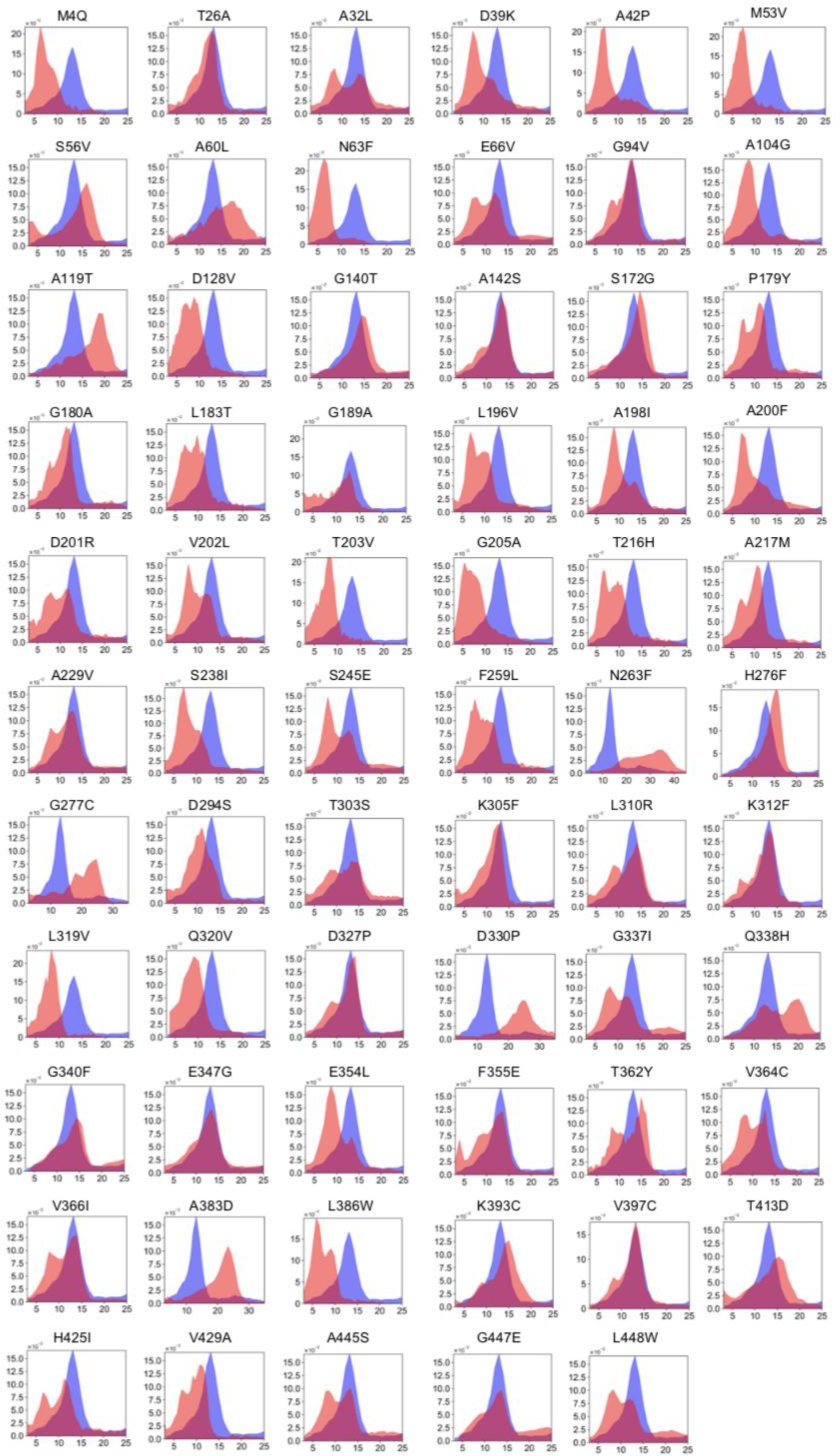
Comparison of distribution of catalytic activities among AP-wt and mutants. X- and y-axes represent catalytic activities (a.u./min) and population percentage of enzyme molecules (peaks 2 and 3). Red and blue bins show the distribution of mutant APs and AP-wt, respectively, except for the highly active mutants.

**Supplementary Figure 8.**
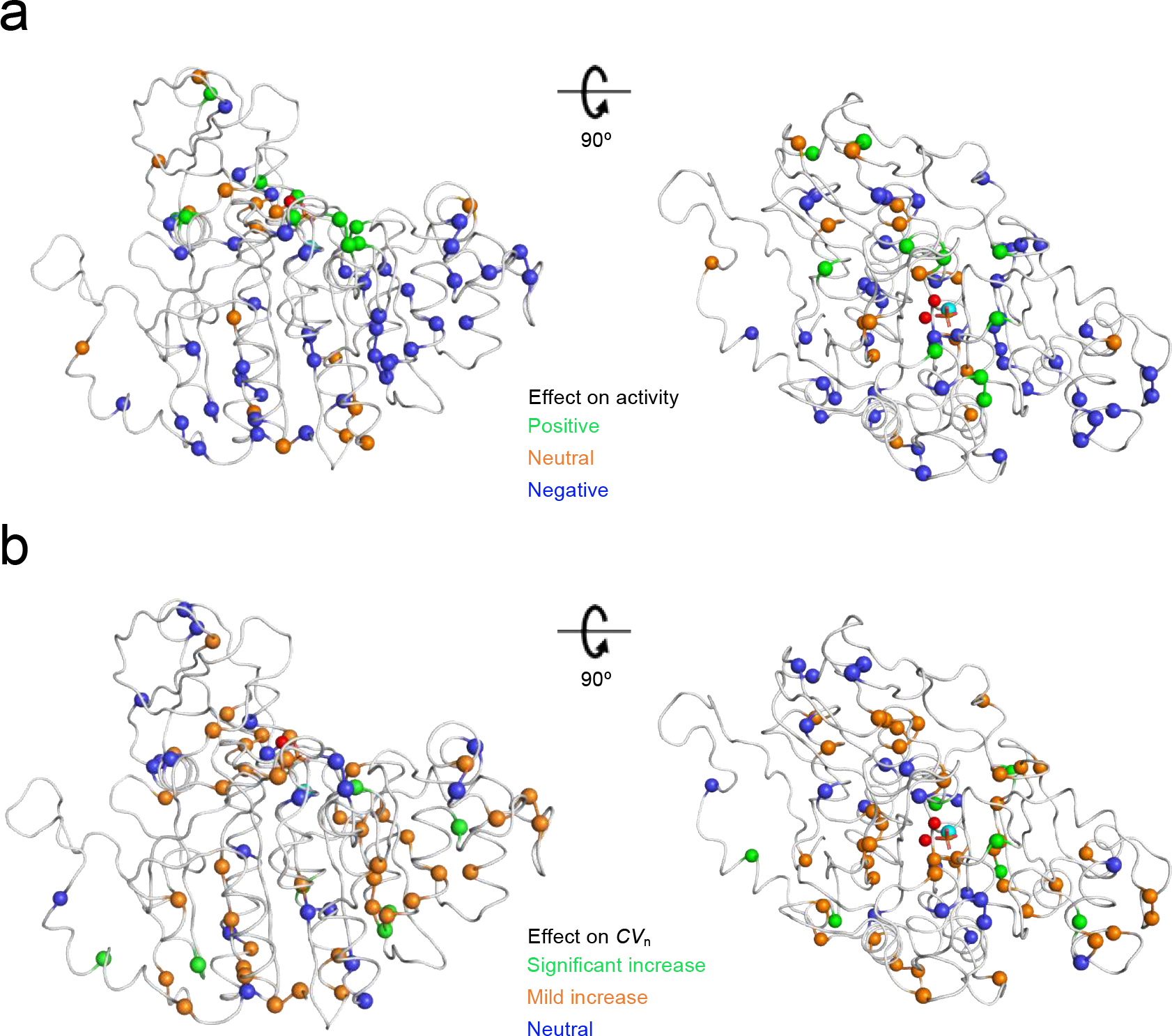
Magnitude of effect of mutation on catalytic activity. **(a) and functional heterogeneity (b) as shown on AP structure.** Spheres represent mutation sites and red and cyan spheres represent zinc and magnesium ions, respectively. The color codes of mean catalytic activity (*MA*) are separated as follows; green (*MA* of mutant > WT + 5 × *SD*), orange (WT – 5 × *SD* < mutant ≤ WT + 5 × *SD*), and blue (mutant ≤ WT – 5 × *SD*). Green, orange, and blue are categorized as positive, neutral, and negative mutations, respectively. The color codes for functional heterogeneity (*CV*_n_) are separated as follows; green (*CV*_n_ of mutant > WT + 5 × *SD*), orange (WT + 2 × *SD* < mutant ≤ WT + 5 × *SD*), and blue (mutant ≤ WT + 2 × *SD*). Green, orange, and blue are categorized as significant increase, mild increase, and neutral, respectively.

**Supplementary Figure 9.**
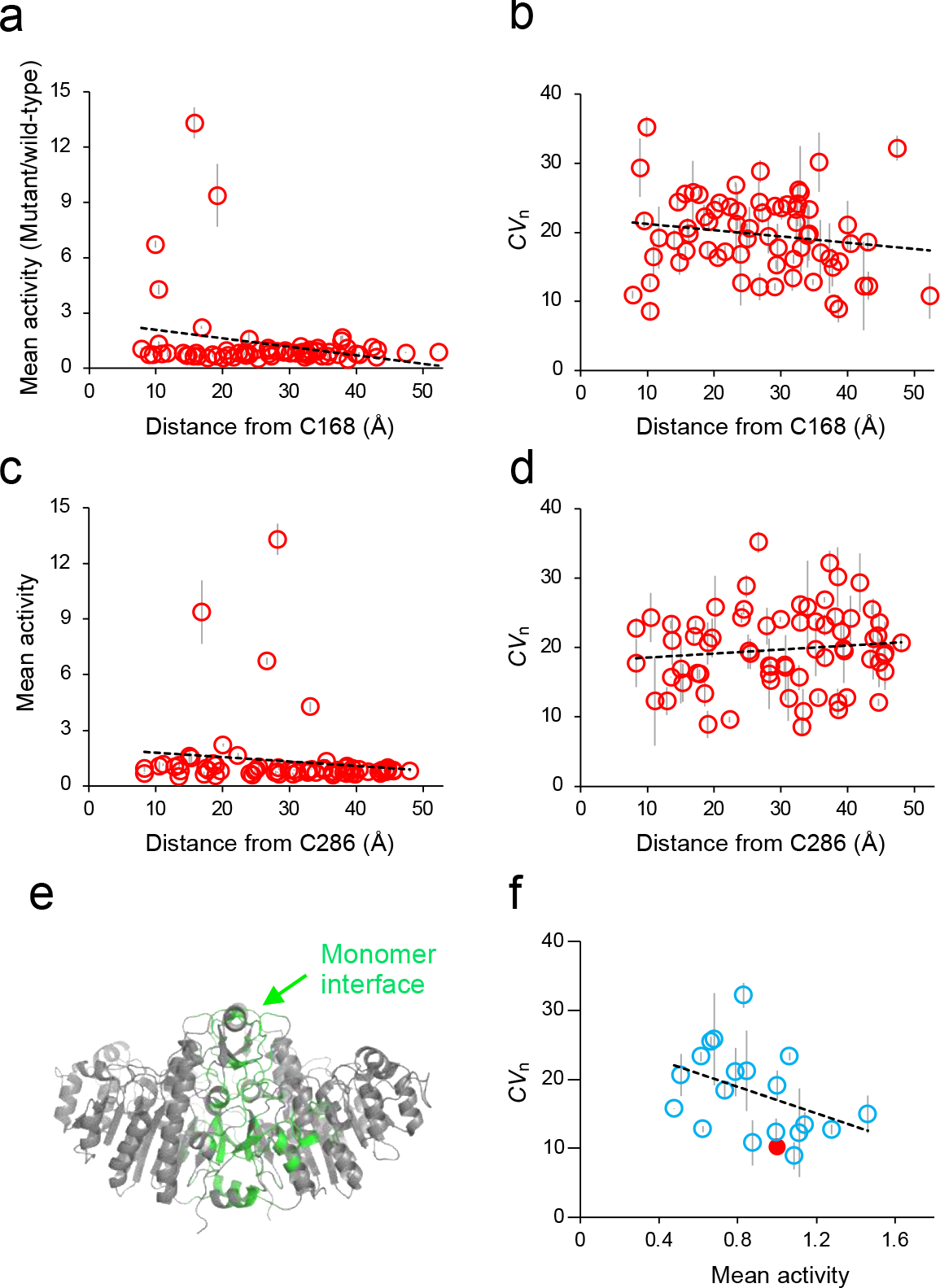
Effects of mutations in catalytic activity and functional heterogeneity on functionally important sites on AP. (**a**) and (**b**) represent the dependence of mean catalytic activity (*MA*) and functional heterogeneity (*CV*_n_) on distance (Å) from C168 (means ± SD). The dotted lines show linear fitting of the plots, and the slopes of (a) and (b) were -0.05 (R^2^ = 0.06) and 0.09 (R^2^ = 0.03), respectively. (**c**) and (**d**) represent the dependence of AP activity and *CV*_n_ on distance (Å) from C286 (means ± *SD*). The slopes of (c) and (d) were -0.02 (R^2^ = 0.02) and 0.06 (R^2^ = 0.01), respectively. (**e**) Sequences located at the interface of the monomers are shown in green. (**f**) Correlation between enzyme activity and functional heterogeneity involving mutations around the monomer interface (mean ± *SD*). The red circle represents AP-wt. The slope of the plot was - 0.10 (R^2^ = 0.17).

## Notes

### Competing Interest Statement

The authors have declared no competing interest.

